# Transient non-integrative nuclear reprogramming promotes multifaceted reversal of aging in human cells

**DOI:** 10.1101/573386

**Authors:** Tapash Jay Sarkar, Marco Quarta, Shravani Mukherjee, Alex Colville, Patrick Paine, Linda Doan, Christopher M. Tran, Constance R. Chu, Steve Horvath, Nidhi Bhutani, Thomas A. Rando, Vittorio Sebastiano

## Abstract

Aging is characterized by a gradual loss of function occurring at the molecular, cellular, tissue and organismal levels^1-3^. At the chromatin level, aging is associated with the progressive accumulation of epigenetic errors that eventually lead to aberrant gene regulation, stem cell exhaustion, senescence, and deregulated cell/tissue homeostasis^3^. The technology of nuclear reprogramming to pluripotency, through over-expression of a small number of transcription factors, can revert both the age and the identity of any cell to that of an embryonic cell by driving epigenetic reprogramming^2,4,5^. Recent evidence has shown that transient transgenic reprogramming can ameliorate age-associated hallmarks and extend lifespan in progeroid mice^6^. However, it is unknown how this form of ‘epigenetic rejuvenation’ would apply to physiologically aged cells and, importantly, how it might translate to human cells. Here we show that transient reprogramming, mediated by transient expression of mRNAs, promotes a rapid reversal of both cellular aging and of epigenetic clock in human fibroblasts and endothelial cells, reduces the inflammatory profile in human chondrocytes, and restores youthful regenerative response to aged, human muscle stem cells, in each case without abolishing cellular identity. Our method, that we named Epigenetic Reprogramming of Aging (ERA), paves the way to a novel, potentially translatable strategy for *ex vivo* cell rejuvenation treatment. In addition, ERA holds promise for *in vivo* tissue rejuvenation therapies to reverse the physiological manifestations of aging and the risk for the development of age-related diseases.

## Main

The process of nuclear reprogramming to Induced Pluripotent Stem cells (iPSCs) is characterized, upon completion, by the resetting of the epigenetic landscape of cells of origin, resulting in reversion of both cellular identity and age to an embryonic-like state ^5,7-9^.

Notably, if the expression of the reprogramming factors is applied only for a short time and then stopped (before the so-called Point of No Return (Nagy and Nagy 2009)), the cells return to the initiating somatic cell state. These observations suggest that if applied for a short enough time (transient reprogramming), the expression of reprogramming factors fails to erase the epigenetic signature defining cell identity; however, it remains unclear whether any substantial and measurable reprogramming of cellular age can be achieved before the PNR and if this can result in any amelioration of cellular function and physiology. To test this, we first evaluated the effect of transient reprogramming on the transcriptome of two distinct cell types - fibroblasts and endothelial cells-from aged human subjects, and we compared it with the transcriptome of the same cell types isolated from young donors (Fig. 1a and e). Fibroblasts were derived from arm and abdomen skin biopsies (25-35 years for the young control, n=3, and 60-70 years for the aged group, n= 3), while endothelial cells were extracted from iliac vein and artery (15-25 years for the young control, n=3, and 45-50 years for the aged group, n=3). We utilized a non-integrative reprogramming protocol that we optimized, based on a cocktail of mRNAs expressing OCT4, SOX2, KLF4, c-MYC, LIN28 and NANOG (OSKMLN)^11^. Our protocol consistently produces induced pluripotent stem cell (iPSC) colonies, regardless of age of the donors, after 12-15 daily transfections; we reasoned that the PNR in our platform occurs at about day 5 of reprogramming, based on the observation that the first detectable expression of endogenous pluripotency-associated lncRNAs occurs at day 5 ^12^. Therefore, we adopted a transient reprogramming protocol where OSKMLN were daily transfected for four consecutive days, and performed gene expression analysis two days after the interruption (Fig. 1b).

**Figure 1.**
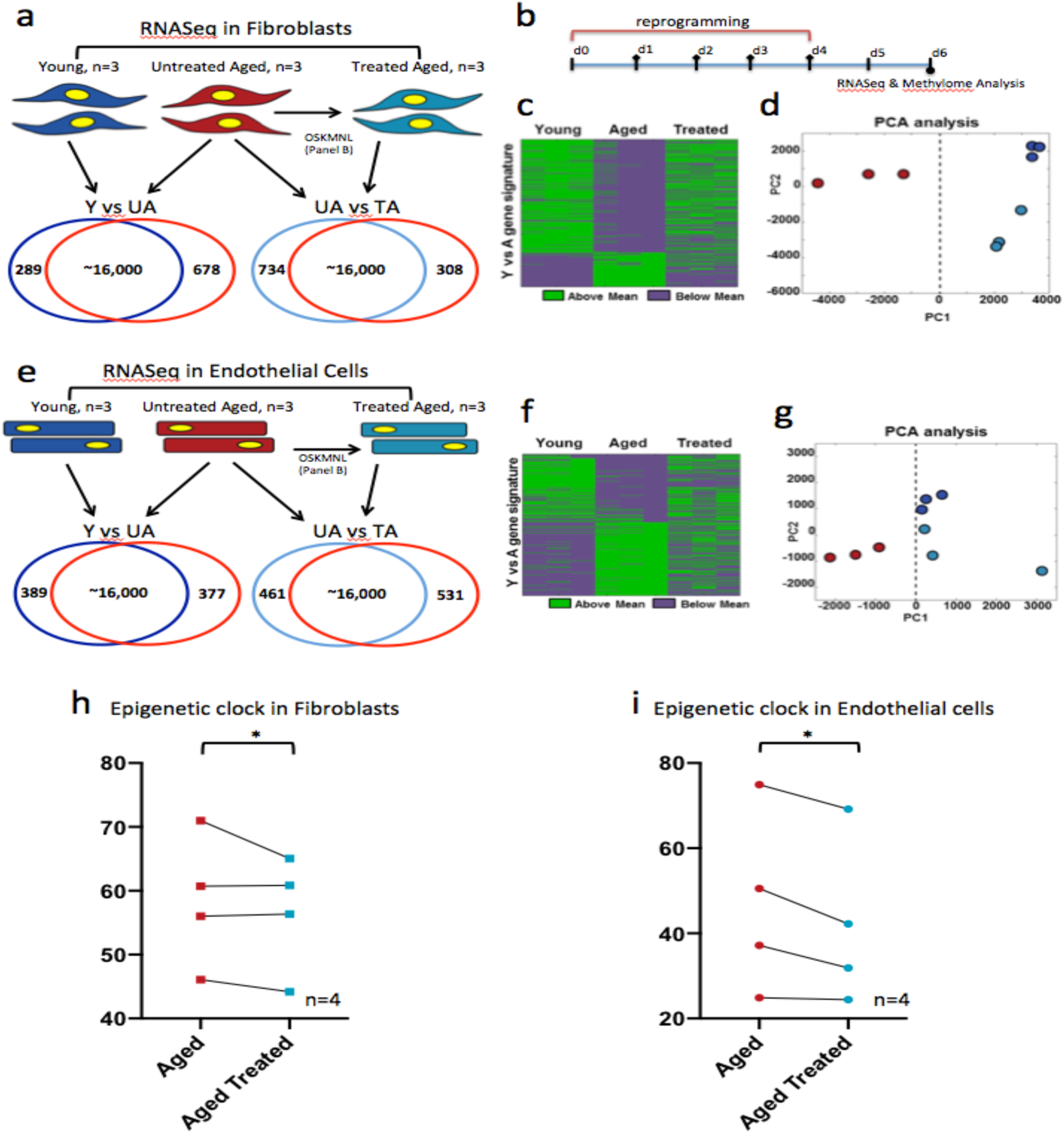
Transcriptomic and Epigenetic clock analysis shows more youthful signature for transiently reprogrammed human fibroblasts and endothelial cells. Cells in each cohort were subjected to 80 bp paired-end reads of RNA sequencing and quantile normalized. a) Venn diagram showing differentially expressed genes defined with a significance p value <0.05 and log fold change > 0.5 between young and aged fibroblasts (group wise comparison) and aged and treated fibroblasts (pairwise comparison). b) Schematic of reprogramming protocol. c) Heat map of polarity of expression (green = above, purple =below) showing the mean for each differential gene. The distribution shows the treated samples transition to a more youthful gene expression pattern in comparison to untreated aged cells D) Principal components analysis in the subspace defined by these differential genes showing clustering of treated and young fibroblasts away from the aged fibroblasts. e) Venn diagram showing differentially expressed genes defined with at significance p value <0.05 and log fold change > 0.5 between young and aged endothelial cells (group wise comparison) and aged and treated endothelial cells (pairwise comparison). f) Heat map of polarity of expression (green = above, purple =below) showing the mean for each differential gene. The distribution shows the treated samples transition to a more expression pattern to the young endothelial cells g) Principal components analysis in the subspace defined by these differential genes showing clustering of treated and young endothelial cells away from the aged endothelial cells. h) Methylation clock estimation of patient sample age with and without treatment for fibroblasts. i) Methylation clock estimation of patient sample age with and without treatment for endothelial cells.

We performed paired-end bulk RNA sequencing on both cell types for the same three cohorts: young (Y), untreated aged (UA), and treated aged (TA). First we compared the quantile normalized transcriptomes of young and untreated aged cells for each cell type (“Y vs UA”) and found that 961 genes (5.85%) in fibroblasts (678 upregulated, 289 downregulated) and 748 genes (4.80%) in endothelial cells (389 upregulated, 377 down regulated) differed between young and aged cells, with the significance criteria of p <.05 and a log fold change cutoff +/-0.5 (Fig. 1a, Supplementary Tables 1, 2). We found these sets of genes were enriched for many of the known aging pathways, identified in the hallmark gene set collection in the Molecular Signatures Database^13^ (Supplementary Table 3, 4). When we mapped the directionality of expression above or below the mean of each gene, we could observe a clear similarity between treated and young cells as opposed to aged cells for both fibroblasts and endothelial cells (Fig. 1c and f). We further performed principal component analysis (PCA) in this gene set space and determined that the young and aged populations were separable along the first principal component (PC1), which explained 64.8% of variance in fibroblasts and 60.9% of variance in endothelial cells. Intriguingly, the treated cells also clustered closer to the younger population along PC1 (Fig. 1d and g).

Using the same significance criteria defined above, we then compared the untreated and treated aged populations (“UA vs TA”) (Fig 1a and e and Supplementary Table 5, 6) and found that 1042 genes in fibroblasts (734 upregulated, 308 downregulated) and 992 in endothelial cells (461 upregulated, 531 downregulated) were differentially expressed. Interestingly, also within these sets of genes we found enrichment for aging pathways, within the Molecular Signatures Database^13^ as previously described (Supplementary Table 7, 8). When we compared the profiles young versus untreated aged (“Y vs UA”) and untreated aged versus treated aged (“UA vs TA”) in each cell type, we observed a 24.7% overlap for fibroblasts (odds ratio of 4.53, p<0.05) and 16.7% overlap for endothelial cells (odds ratio of 3.84, p<0.05) with the directionality of change in gene expression matching that of youth (i.e. if higher in young then higher in treated aged); less than 0.5% moved oppositely in either cell types (Fig. S1, Supplementary Table 9, 10).

Next, we used these transcriptomic profiles to verify retention of cell identity after transient reprogramming. To this end, using established cell identity markers, we verified that none significantly changed upon treatment (Supplementary Table 11). In addition, we could not detect the expression of any pluripotency-associated markers (other than the OSKMLN mRNAs transfected in) (Supplementary Table 11). Altogether, the analysis of the transcriptomic signatures revealed that transient reprogramming triggers a more youthful gene expression profile, while retaining cell identity.

Epigenetic clocks based on DNA methylation levels are the most accurate molecular biomarkers of age across tissues and cell types and are predictive of a host of age-related conditions including lifespan ^5,14-16^. Exogenous expression of canonical reprogramming factors (OSKM) is known to revert the epigenetic age of primary cells to a prenatal state^5^. To test whether transient expression of OSKMLN could reverse the epigenetic clock, we used two epigenetic clocks that apply to human fibroblasts and endothelial cells: Horvath’s original pan-tissue epigenetic clock (based on 353 cytosine-phosphate-guanine pairs), and the more recent skin & blood clock (based on 391 CpGs)^5,17^.

According to the pan tissue epigenetic clock, transient OSKMLN significantly (two-sided mixed effects model P-value=0.023) reverted the DNA methylation age (average age difference=-3.40 years, standard error 1.17). The rejuvenation effect was more pronounced in endothelial cells (average age difference=-4.94 years, SE=1.63, Figure 1i) than in fibroblasts (average age difference=-1.84, SE=1.46, Figure 1h). Qualitatively similar, but less significant results, could be obtained with the skin and blood epigenetic clock (overall rejuvenation effect −1.35 years, SE=0.67, one-sided mixed effects model P-value=0.042, average rejuvenation in endothelial cells and fibroblasts is −1.62 years and −1.07, respectively).

Prompted by these results, we next analyzed the effect of transient reprogramming on various hallmarks of cellular physiological aging. We employed a panel of 11 established assays, spanning the hallmarks of aging^18^ (Supplementary Table 12), and performed most of the analyses using single cell high throughput imaging to capture quantitative changes in single cells and distribution shifts in the entire population of cells. All the analyses were performed separately in each individual cell line (total of 19 fibroblast lines: 3 young, 8 aged and 8 treated aged; total 17 endothelial cell lines: 3 young, 7 aged and 7 treated aged) (Fig. 2a, Figures S2-5). Statistical analysis was conducted in each paired sets of samples; the data was subsequently pooled by age category for ease of representation (Fig. 2a, see Materials and Methods for a detailed description of the Statistical methods that were used). Control experiments were performed by adopting the same transfection scheme using mRNA encoding for GFP (Figures S6, 7).

**Figure 2.**
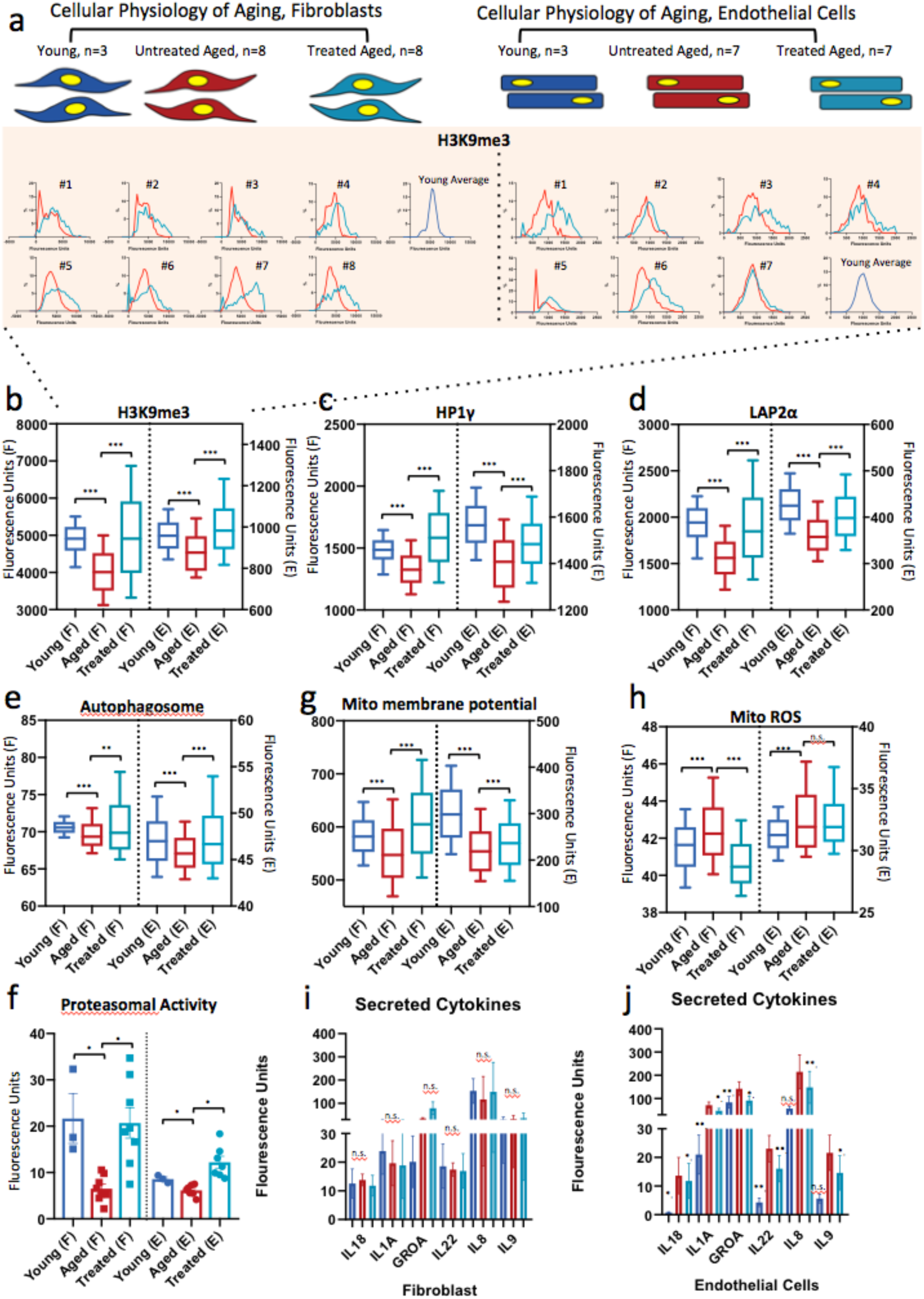
Transient reprogramming reverts aged physiology towards a more youthful state in human fibroblasts and endothelial cells: a) Workflow summarizing the strategy adopted to assess reversal of physiological aging in cells. Fibroblasts and endothelial cells were obtained from otherwise healthy young and aged individuals. Young untreated cells (dark blue), aged untreated cells (red), and transiently reprogrammed aged cells (light blue) were analyzed for a panel of 11 different hallmarks of aging. Most of the assays were performed by high throughput imaging to allow population-wide studies with single cell resolution (in the example H3K9me3). Distribution data was collected for each patient in each condition. However for simplicity in depiction, the fluorescent baseline of each was adjusted so that the mean of each aged control was the same, with the corresponding normalization of the other conditions. This allowed the distributions to align, be overlayed and depicted once per condition: i.e. young, aged and treated, as shown in the example histogram on the right side of panel a. These distributions were depicted as boxplots for further simplicity in depiction (on the left side of the dotted line fibroblasts; on the right side of the dotted line endothelial cells). T-tests p values were calculated before this normalization comparing the pooled young and with three of the aged distributions evaluated during the same run. While p values for treatment effect were calculated comparing the individual distributions of treated and corresponding untreated patient cells (in which case the overall significance ranking was set by the second most stringent of the p values, to allow one grace for patient to patient variability). P value: *<.05, **<.01, ***<.001 b) Quantification of single nucleus levels of trimethlyated H3K9, which binds to HP1 to constitutive heterochomatin and is considered a repressive mark of gene expression^10^. Both cell types show significant elevation of the mark towards the youthful distribution. c) Quantification of single nucleus levels of heterochromatin marker HP1γ itself by immunocytochemistry again showing a trend toward youth after cells are treated. d) Quantification of the inner nuclear membrane polypeptide LAP2α which regulates nuclear lamina by regulating the binding of lamin B1 and chromatin. This again shows a trend toward youth after cells are treated. e) Results of live cells imaging with florescent marker of autophagosome formation in single cells. Simply after two days of relaxation, the treated distribution moves above even the youthful one, indicative of the wave of protein clearing engaged by early reprograming. f) Cleavage of fluorescent-tagged chymotrypsin like substrate elevated in treated and young fibroblasts and endothelial cells corresponding to increased proteasome 20S core particle activity. g) Individual cell mitochondria membrane potential measurements also showing more active mitochondria as a result of transient reprogramming. h) Individual cell mitochondria ROS measurements also showing less accumulated ROS as a result of transient reprogramming. i) Quantification of pro-inflammatory factors secreted by fibroblasts. j) Quantification of pro-inflammatory factors secreted by endothelial cells. The levels of the these factors were significantly elevated in the aged endothelial cohorts and subsequently diminished after treatment, using a p<.05 cutoff and fold change higher than 1.5,. Error bars represent standard deviation of the mean.

To extend our previous findings on epigenetics, we quantitatively measured by immunofluorescence (IF) the epigenetic repressive mark H3K9me3, the heterochromatin-associated protein HP1**γ**, and the nuclear lamina support protein LAP2α (Fig 2b-d). Aged fibroblasts and endothelial cells showed a decrease in the nuclear signal for all three markers compared to young cells, as previously reported^8^; treatment of aged cells resulted in an increase of these markers in both cell types. Next, we examined both pathways involved in proteolytic activity of the cells by measuring formation of autophagosomes, and chymotrypsin-like proteasomal activity, both reported to decrease with age ^19,20^. Treatment increased both to levels similar to or even higher than young cells, suggesting that early steps in reprogramming promote an active clearance of degraded biomolecules (Fig. 2 e, f).

In terms of energy metabolism, aged cells display decreased mitochondrial activity, accumulation of reactive oxygen species (ROS), and deregulated nutrient sensing^8,20,21^. We therefore tested the effects of treatment on aged cells by measuring mitochondria membrane potential, mitochondrial ROS, and levels of Sirtuin1 protein (SIRT1) in the cells. Transient reprogramming increased mitochondria membrane potential in both cell types (Fig. 2g), while it decreased ROS (Fig. 2h) and increased SIRT1 protein levels in fibroblasts, similar to young cells (Figure S8). Senescence-associated beta-galactosidase staining showed a significant reduction in the number of senescent cells in aged endothelial cells (Figure S8). This decrease was accompanied by a decrease in pro-inflammatory senescence associated secretory phenotype (SASP) cytokines (Fig 1) again in endothelial cells. ^20,22,23^. Lastly, in neither cell type did telomere length, measured by quantitative fluorescence in situ hybridization ^8,24^, show significant extension with treatment (Fig. S8), suggesting that the cells did not de-differentiate into a stem-like state in which telomerase activity would be reactivated.

Next, we assessed the perdurance of these effects and found that most were significantly retained after four and six days from the interruption of reprogramming (Figs S9,10). We then examined how rapidly these physiological rejuvenative changes manifest by repeating the same sets of experiments in fibroblasts and endothelial cells that were transfected for just two consecutive days. Remarkably, we observed that most of the rejuvenative effects could already be seen after two days of treatment, although most were more moderate (Figs S11, 12).

Collectively, this data demonstrate that transient expression of OSKMLN can induce a rapid, persistent reversal of cellular age in human cells at the transcriptomic, epigenetic and cellular levels. Importantly, these data demonstrate that the process of “cellular rejuvenation” - that we name Epigenetic Reprogramming of Aging, or “ERA” - is engaged very early and rapidly in the iPSC reprogramming process. These epigenetic and transcriptional changes occur before any epigenetic reprogramming of cellular identity takes place, a novel finding in the field.

With these indications of a beneficial effect of ERA on cellular aging, we next investigated whether ERA could also reverse the inflammatory phenotypes associated with aging. After obtaining preliminary evidence of this reversal in endothelial cells (Figure 2i, j), we extended our analysis to osteoarthritis, a disease strongly associated with aging and characterized by a pronounced inflammatory spectrum affecting the chondrocytes within the joint^25^. We thus isolated chondrocytes from cartilage of 60-70 year old patients undergoing total joint replacement surgery owing to their advanced stage OA and compared the results of treatment to chondrocytes isolated from young individuals (Fig. 3a). Transient reprogramming was performed for two or three days and the analysis performed after two days from interruption of reprogramming, though the more consistent effect across patients was with longer treatment. Treatment showed a significant reduction in pro-inflammatory cytokines (Fig. 3b), intracellular mRNA levels of RANKL and iNOS2, as well as in levels of inflammatory factors secreted by the cells (Fig. 3 b-d). In addition, ERA promoted cell proliferation (Fig. 3e), increased ATP production (Fig. 3f), and decreased oxidative stress as revealed by reduced mitochondrial ROS and elevated RNA levels of antioxidant SOD2 (Fig. 3g, h), a gene that has been shown to be downregulated in OA^26^., We observed that ERA did not affect the expression level of SOX9 (a transcription factor core to chondrocyte identity and function) and significantly increased the level of expression of COL2A1 (the primary collagen in articular cartilage) (qRT-PCR in Fig 3i, j), suggesting retention of chrondrogenic cell identity. Together, these results show that transient expression of OSKMLN can promote a partial reversal of gene expression and cellular physiology in aged OA chondrocytes toward a healthier state, suggesting a potential new therapeutic strategy to ameliorate the OA disease process.

**Figure 3.**
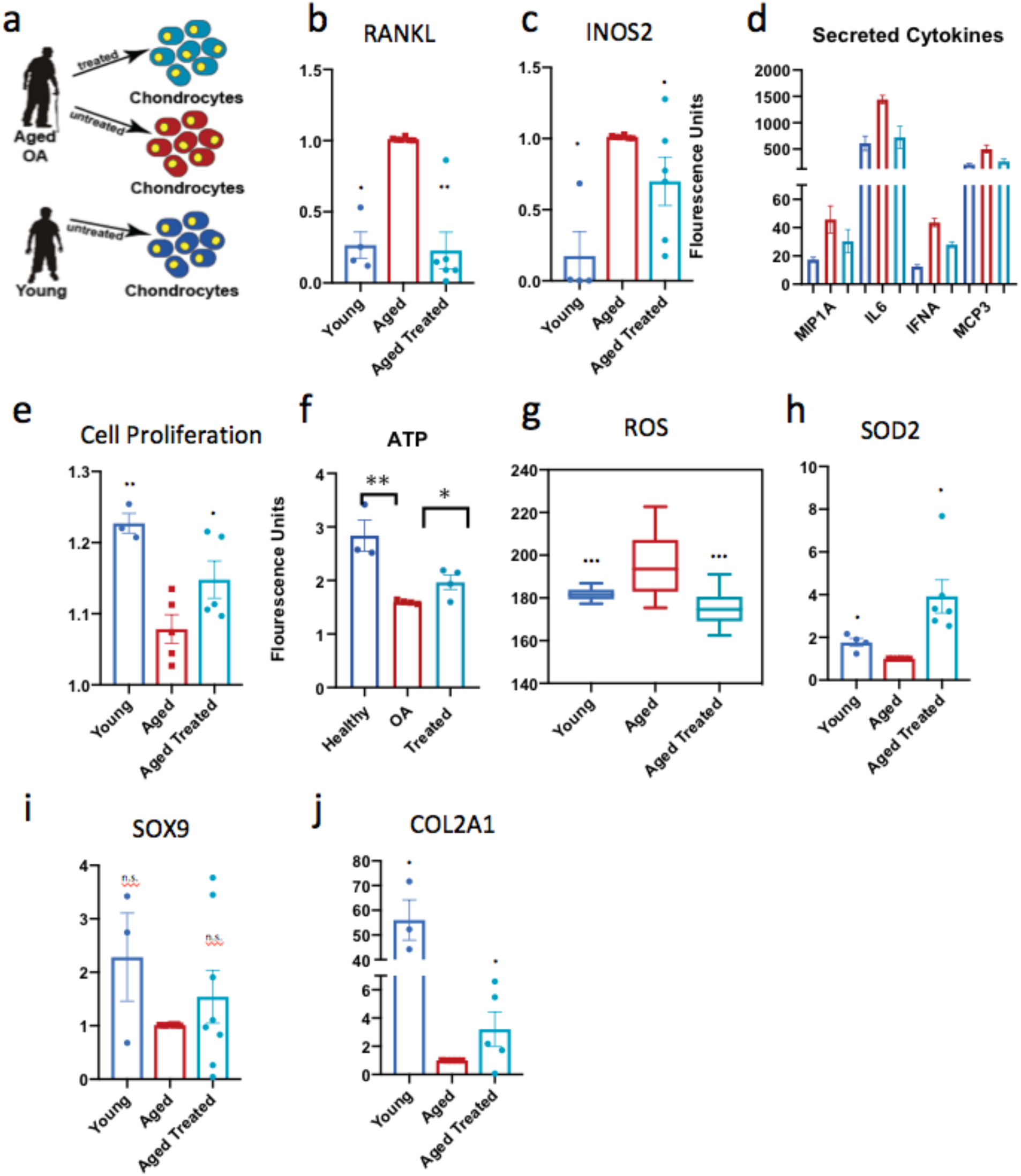
Transient reprogramming mitigates inflammatory phenotypes in diseased chondrocytes. a) Workflow summarizing the strategy adopted to assess amelioration of age related disease. Chondrocytes were obtained from young (blue) and aged diagnosed late stage Osteoarthritis (OA) patients from cartilage biopsies. Aged OA cells (red) and transiently reprogrammed OA cells (light blue) were evaluated for OA specific phenotypes. All box and bar plots are combined over biological and technical replicates for ease of viewing and compared as in Fig 1 –significance ranking was set by second most stringent p value. Significance is calculated with students t-tests P value: *<.05, **<.01, ***<.001. Error bars show s.d. b) qRT-PCR evaluation of RNA levels of RANKL. c) qRT-PCR evaluation of RNA levels of INOS2. d) Cytokine profiling of chondrocyte secretions shows an increase pro-inflammatory cytokines that diminishes with treatment. E) Cell proliferation. f) Measurement of ATP concentration using glycerol based fluorophore shows elevation of ATP levels with treatment. g) Individual cell mitochondria ROS measurements also showing less accumulated ROS as a result of transient reprogramming. h) qRT-PCR evaluation of RNA levels of antioxidant SOD2. i) qRT-PCR levels of chondrogenic identity and function transcription factor SOX9 is retained after treatment. f) qRT-PCR shows elevation RNA levels for extracellular matrix protein component COL2A1.

Stem cell loss of function and regenerative capacity represents another important hallmark of aging^18^ We thus wanted to assess the effect of transient reprogramming on the age-related changes in somatic stem cells that impair regeneration. First, we tested the effect of transient reprogramming on mouse-derived skeletal muscle stem cells (MuSCs). We treated MuSCs for 2 days while they were kept in a quiescent state using an artificial niche^27^. We conducted initial experiments with young (3 month) and aged (20-24 months) murine MuSCs isolated by FACS (Fig. 4a). Treatment of aged MuSCs reduced both time of first division, approaching the faster activation kinetics of quiescent young MuSCs^28,29^, and mitochondrial mass ^30^ (Fig. S13a-b). Moreover, treatment partially rescued the reduced ability of single MuSCs to form colonies^28,31^ (Fig. S13c). We further cultured these cells and observed that treatment did not change expression of the myogenic marker MyoD but instead improved their capacity to differentiate into myotubes (Fig. S14a-d), suggesting that transient reprogramming does not disrupt the myogenic fate but can enhance the myogenic potential.

**Figure 4.**
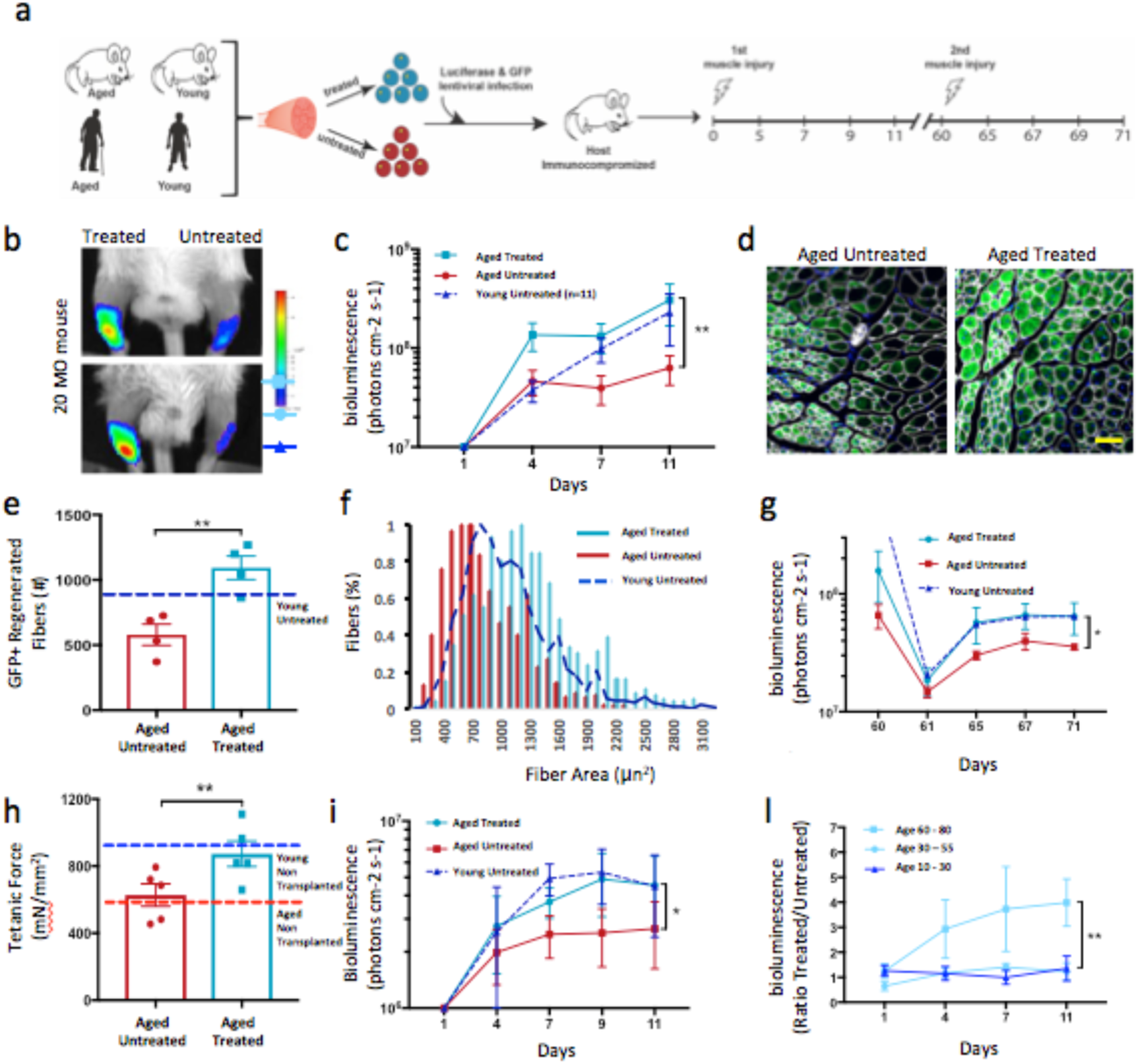
Transient Reprogramming restores aged Muscle Stem Cell potency. a) Schematic showing the experimental design of partially reprogrammed aged mouse and human MuSCs b) Representative images of bioluminescence measured from mice 11 days after transplantation and injury in TA muscles of treated/untreated Luciferase^+^ mouse MuSCs c) Quantified results of bioluminescence in (b) at different time points following transplantation and injury (n=10). d) Representative immunofluorescence of GFP expression in TA muscle cross-sections of mice imaged and quantified in c and d, isolated 11 days after transplantation (Error bar = 500 μm). e) Quantification of immunofluorescence staining in (d) (n=5). f) Quantification of cross sectional area of donor derived GFP+ fibers in TA muscles that were recipients of transplanted MuSCs (n=5). g) Results of bioluminescence imaging of TA muscles reinjured after 60 days (second injury) after MuSC transplantations (n=6). The second injury was performed to test whether the bioluminescence signal increased as a consequence of activating and expanding luciferase^+^/GFP^+^ MuSCs that were initially transplanted and that had engrafted under the basal lamina. h) Tetanic force measurements of aged muscles injured and transplanted with aged MuSCs. TA muscles were dissected and electrophysiology ex vivo for tetanic measurement performed. Baseline of force production of untransplanted muscles was measured in young (4 months, blue broken line) and aged (27 months, red broken line) mice. Treated aged MuSCs were transplanted into TA muscles of aged mice and force production measured 30 days later (n=5). i) Quantified results of bioluminescence measured from mice 11 days after transplantation in TA muscles of treated Luciferase^+^ human MuSCs. l) Variation in ratio of bioluminescence between treated and untreated MuSCs obtained from healthy donors of different age groups. Significance is calculated with students t-test, pairwise between treated and aged, and group wise when comparing to young patients (Age groups. 10-30: n=5; 30-55: n=7; 60-80: n=5). P value: *<.05, **<.01, ***<.001 color of asterisks match population being compared to.

Next, we wanted to test MuSC function and potency to regenerate new tissue *in vivo*. To do this, we transduced young, aged, or transiently reprogrammed aged MuSCs with a lentivirus expressing luciferase and green fluorescent protein (GFP) and then transplanted the cells into injured tibialis anterior (TA) muscles of immunocompromised mice. Longitudinal bioluminescence imaging (BLI) initially showed that muscles transplanted with treated aged MuSCs showed the highest signal (day 4, Fig 4b, c), but became comparable to muscles with young MuSCs by day 11 post-transplantation; conversely muscle with untreated aged MuSCs showed lower signals at all time points post transplantation (Figure 4b, c). Immunofluorescence analysis further revealed higher numbers of donor-derived (GFP^+^) myofibers in TAs transplanted with treated compared to untreated aged MuSCs (Fig. 4d, e). Moreover, the GFP^+^ myofibers from treated aged cells exhibited increased cross-sectional areas when compared to their untreated counterparts, and in fact even larger than the young controls (Figure 4f). Together, these results suggest improved tissue regenerative potential of transiently reprogrammed aged MuSCs. After 3 months, all mice were subjected to autopsy, and no neoplastic lesions or teratomas were discovered (Supplementary Table 13). To test potential long-term benefits of the treatment, we induced a second injury 60 days after cell transplantation, and again observed that TA muscles transplanted with transiently reprogrammed aged MuSCs yielded higher BLI signals (Figure 4g).

Sarcopenia is an age-related condition that is characterized by loss of muscle mass and force production^32,33^. Similarly, in mice muscle functions show progressive degeneration with age^34,35^.We wanted to test whether transient reprogramming of aged MuSCs would improve a cell-based treatment in restoring physiological functions of muscle of older mice. To test this, we first performed electrophysiology to measure tetanic force production in TA muscles isolated from young (4 months) or aged (27 months) immunocompromised mice. We found that TA muscles from aged mice have lower tetanic forces compared to young mice, suggesting an age-related loss of force production (Fig. 4h). Next, we isolated MuSCs from aged mice (20-24 months). After treating aged MuSCs, we transplanted them into cardiotoxin-injured TA muscles of aged (27 months) immunocompromised mice. We waited 30 days to give enough time to the transplanted muscles to fully regenerate. We then performed electrophysiology to measure tetanic force production. Muscles transplanted with untreated aged MuSCs showed forces comparable to untransplanted muscles from aged control mice (Fig. 4h). Conversely, muscles that received treated aged MuSCs showed tetanic forces comparable to untransplanted muscles from young control mice (Fig. 4h and S15a). These results suggest that transient reprogramming in combination with MuSC-based therapy can restore physiological function of aged muscles to that of youthful muscles.

Lastly, we wanted to translate these results to human MuSCs. We repeated the study, employing operative samples obtained from patients in different age ranges (10 to 80 years old), and transducing them with GFP- and luciferase-expressing lentiviral vectors (Fig. 4a). As in mice, transplanted, transiently-reprogrammed, aged human MuSCs resulted in increased BLI signals compared to untreated MuSCs from the same individual and comparable to those observed with young MuSCs (Figure 4I and S16a-b). Interestingly, the BLI signal ratio between contralateral muscles with treated and untreated MuSCs was higher in the older age group (60 – 80 year old) than in the younger age groups (10-30 or 30-55 years old), suggesting that ERA restores lost functions to younger levels in aged cells (Figure 4l). Taken together, these results suggest that transient reprogramming partially restores the potency of aged MuSCs to a degree similar to that of young MuSCs, without compromising their fate, and thus has potential as a cell therapy in regenerative medicine.

Nuclear reprogramming to induced pluripotent stem cells (iPSCs) is a multi-phased process comprising initiation, maturation and stabilization^36^. Upon completion of such a dynamic and complex “epigenetic reprogramming”, iPSCs are not only pluripotent but also youthful. Here we have demonstrated that a non-integrative, mRNAs-based platform of transient cellular reprogramming can very rapidly reverse a broad spectrum of aging hallmarks in the initiation phase, when epigenetic erasure of cell identity has not yet occurred. We show that the process of rejuvenation occurs in naturally aged human cells, with restoration of lost functionality in diseased cells and aged stem cells while preserving cellular identify. Future studies are required to elucidate the mechanism that drives the reversal of the aged phenotype during cellular reprogramming, uncoupling it from dedifferentiation process^37,38^. While proof of principle of this was shown in a genetic model of aging (progeroid mice) the proof that a multispectral cellular rejuvenation could be achieved in human cells isolated from naturally aged individuals was missing. Our results are novel and represent a significant step toward the goal of reversing cellular aging and have potential therapeutic implications for aging and aging-related diseases.

## Supporting information

UY vs UA gene signature fibroblasts

UY vs UA gene signature endothelial cells

MSigDB UY vs UA fibdrobalsts.

MSigDB UY vs UA endothelial cells

TA vs UA gene signature fibroblasts

TA vs UA gene signature endothelial cells.

MSigDB UA vs TA endothelial cells

MSigDB UA vs TA fibroblasts

overlap signatures fibroblasts

overlap signatures endothelial cells

## Acknowledgments

We thank the members of the Sebastiano, Bhutani and Rando laboratories for comments and discussions; we thank Jens Durruthy Durruthy for valuable technical information on mRNA-based transient reprogramming. We thank Tony Wyss-Coray and his lab for providing primary fibroblast lines from patients diagnosed with AD and Helen Blau and her lab for technical consulting on Telomere length evaluation. This work was supported by the AFAR Junior Investigator Award to V.S., by grants from the National Institutes of Health (NIH/NIAMS) (R01 AR070865 and R01 AR070864) to N.B., the Glenn Foundation for Medical Research and by grants from the National Institutes of Health (NIH) (P01 AG036695, R01 AG23806 (R37 MERIT Award), R01 AG057433, and R01 AG047820), and the Department of Veterans Affairs (BLR&D and RR&D Merit Reviews) to T.A.R.; and by the CalPoly funding Award # TB1-01175.

## Author Contributions

V.S., T.A.R., T.J.S., N.B., and M.Q. conceived, designed and supervised the experiments reported. T.J.S. performed the Fibroblast and Endothelial *in vitro* experiments. T.J.S. performed the processing for the Fibroblast and Endothelial RNA sequencing experiments. N.B. and C.C. obtained the cartilage surgical specimens and S.M and T.J.S performed the chondrocyte isolation and *in vitro* experiments. P.P., C.M.T. and A.C. performed the MuSCs isolation, and lentivirus infection design and experiments. T.J.S., P.P., C.M.T. and A.C. performed the MuSCs *in vitro* experiments. M.Q., P.P., A.C. and L.D. performed the MuSCs *in vivo* experiments. T.J.S., S.M, N.B., M.Q., C.C., S.H., T.A.R., and V.S. analyzed data and wrote the manuscript.

## DATA AVAILABILITY

The data that support the findings of this study are available from the corresponding author upon request.

## Methods

### Human Fibroblast Isolation and Culture

Isolation was performed at Coriell Institute on healthy patients and from Alzheimer patient samples at Stanford Hospital, in accordance to the methods and protocols approved by Institutional Review Board of Stanford University, biopsied for skin mesial aspect of mid-upper arm or abdomen using 2 mm-punch biopsies from both male and female patients 60-70 years old (n=8) and 25-35 years (n=3). Cells were cultured out from these explants and maintained in Eagle’s Minimum Essential Medium with Earl’s salts supplemented with non-essential amino acids, 10% fetal bovine serum and 1% Penicillin/Streptomycin. Cells were cultured at 37°C with 5% CO2.

### Human Endothelial Cell Isolation and Culture

Isolation was performed at Coriell Institute from iliac arteries and veins and muscle biopsies from Stanford Hospital, in accordance to the methods and protocols approved by Institutional Review Board of Stanford University, from otherwise healthy 45-60 years old (n=7). Tissue was digested with collagenase and cells released from the lumen were used to initiate cultures. Plates for seeding were coated with 2% gelatin, then washed with PBS before use. Cells were maintained in Medium 199 supplemented with 2 mM L-glutamine, 15% fetal bovine serum, 0.02 mg/ml Endothelial Growth Supplement, 0.05 mg/ml Heparin and 1% Penicillin/Streptomycin. Cells were cultured at 37°C with 5% CO2.

### Human Articular Chondrocyte Isolation and Culture

In accordance to the methods and protocols approved by Institutional Review Board of Stanford University, the human OA chondrocytes were derived from discarded tissues of OA patients (50-72 years of age, n=6) undergoing total knee arthroplasty. The samples were surgical waste and were fully deidentified prior to procurement, hence no prior patient consent was required. Cartilage pieces were shaved off bone by scalpel, taking care to avoid any fat, then digested with collagenase in DMEM/F12 media (supplemented with 25mg/ml ascorbate, 2mM L-glutamine, 1% penicillin/streptomycin antibiotics and 10% fetal bovine serum) for one to two days until shavings were substantially dissolved. Supernatant from cultures was strained, filtered and centrifuged and the cells were then resuspended in fresh media. The chondrocytes were cultured in high density monolayer in at 37°C with 5% CO2.

### Mice

C57BL/6 male mice and NSG mice were obtained from Jackson Laboratory. NOD/MrkBomTac-Prkdcscid mice were obtained from Taconic Biosciences. Mice were housed and maintained in the Veterinary Medical Unit at the Veterans Affairs Palo Alto Health Care Systems. The Administrative Panel on Laboratory Animal Care of Stanford University approved animal protocols.

### Human skeletal muscle specimens

The human muscle biopsy specimens were taken after patients (10-30 years, n=2; 30-55 years, n=2; 60-80 years, n= 3) gave informed consent as part of a human studies research protocol that was approved by the Stanford University Institutional Review Board. Sample processing for cell analysis began within one to twelve hours of specimen isolation. In all studies, standard deviation reflects variability in data derived from studies using true biological replicates (that is, unique donors). Data were not correlated with donor identity.

### MuSC Isolation and Purification

Muscles were harvested from mouse hind limbs (n=4) and mechanically dissociated to yield a fragmented muscle suspension. This was followed by a 45-50 minute digestion in a Collagenase II-Ham’s F10 solution (500 units per ml; Invitrogen). After washing, a second digestion was performed for 30 minutes with Collagenase II (100 units per ml) and Dispase (2 units per ml; ThermoFisher). The resulting cell suspension was washed, filtered and stained with VCAM-biotin, CD31-FITC, CD45-APC and Sca-1-Pacific-Blue antibodies, all at dilutions of 1:100. Human MuSCs were purified from fresh operative samples^50,51^. Operative samples were carefully dissected from adipose and fibrotic tissue and a dissociated muscle suspension prepared as described for mouse tissue. The resulting cell suspension was then washed, filtered and stained with anti-CD31-Alexa Fluor 488, anti-CD45-Alexa Fluor 488, anti-CD34-FITC, anti-CD29-APC and anti-NCAM-Biotin antibodies. Unconjugated primary antibodies were then washed and the cells were incubated for 15 min at 4°C in streptavidin-PE/Cy7 to detect NCAM-biotin. Cell sorting was performed on calibrated BD-FACS Aria II or BD FACSAria III flow cytometers equipped with 488-nm, 633-nm and 405-nm lasers to obtain the MuSC population. A small fraction of sorted cells was plated and stained for Pax7 and MyoD to assess the purity of the sorted population.

### mRNA Transfection

Cells were transfected using either mRNA-In (mTI Global Stem) for fibroblasts and chondrocytes, to reduce cell toxicity, and Lipofectamine MessengerMax (Thermo Fisher) for endothelial cells and MuSCs, which were more difficult to transfect, using manufacturer’s protocol. For fibroblast and endothelial cells, serum free Pluriton medium with bFGF was used for transfection, while muscle stem cells and chondrocytes were kept in their original media – the former lacking serum and the later requiring serum to prevent the natural dedifferentiation of chondrocytes in culture. Culture medium was changed for fibroblasts and endothelial cells 4 hours after transfection, but not for chondrocytes or MuSCs as overnight incubation was needed to produce a significant uptake of mRNA. Efficiency of delivery was confirmed by both GFP mRNA and immunostaining for individual factors in OSKMLN cocktail, the former also being used as a transfection control with the same protocol.

### Immunocytochemistry

Cells were washed with HBSS/CA/MG then fixed with 15% paraformaldehyde in PBS for 15 minutes. Cells were then blocked for 30 min to one hour with a blocking solution of 1% BSA and 0.3% Triton X-100 in PBS for fibroblasts, endothelial cells and 20% donkey serum/0.3% Triton in PBS for MuSCs. Primary antibodies were then applied in blocking solution and allowed to incubate overnight at 4°C. The following day, the cells were washed with HBSS/CA/MGor PBST for MuSCs before switching to the corresponding Alexa Flour-labeled secondary antibodies and incubated for 2 hours. The cells were then washed again and stained with DAPI for 30 minutes and switched to HBSS/CA/MGfor imaging or Fluoview for MuSCs.

### Autophagosome Formation Staining

Cells were washed with HBSS/Ca/Mg and switched to a staining solution containing a proprietary fluorescent autophagosome marker (Sigma). The cells were then incubated at 37°C in 5% CO_2_ for 20 minutes, washed 2 times using HBSS/Ca/Mg, and stained for 15 minutes using CellTracker Deep Red cell labeling dye. Cells were then switched to HBSS/Ca/Mg for single cell imaging using the Operetta High Content Imaging System (Perkin Elmer)

### Proteasome Activity Measurement

Wells were first stained with PrestoBlue Cell Viability dye (Life Technologies) for 10 minutes. Well signals were read using a TECAN fluorescent plate reader as a measure of cell count. Then cells were washed with HBSS/Ca/Mg before switching to original media containing the chymotrypsin like fluorogenic substrate LLVY-R110 (Sigma), which is cleaved by proteasome 20S core particle. Cells were then incubated at 37°C in 5% CO_2_ for 2 hours before signals were again read on the TECAN fluorescent plate reader. Readings were then normalized by PrestoBlue cell count.

### Mitochondrial Membrane Potential Staining

Tetramethylrhodamine Methyl Ester Perchlorate (Thermo) was added to cell culture media. This dye is sequestered by active mitochondria based on their membrane potential. Cells were incubated for 30 minutes at 37°C in 5% CO_2_ and washed 2 times with HBSS/Ca/Mg before staining for 15 minutes using CellTracker Deep Red. Finally, cells were imaged in fresh HBSS/Ca/Mg using the Operetta High Content Imaging System (Perkin Elmer)

### Mitochondrial ROS Measurement

Cells were washed with HBSS/Ca/Mg and then switched to HBSS/Ca/Mg containing MitoSOX (Thermo), a live cell permeant flurogenic dye that selectively targeted to mitochondria and fluoresces when oxidized by superoxide. Cells were incubated for 10 minutes at 37°C in 5% CO_2_. Cells were then washed twice with HBSS/Ca/Mg, and stained for 15 minutes using CellTracker Deep Red. Finally, cells were imaged in fresh HBSS/Ca/Mg using the Operetta High Content Imaging System (Perkin Elmer)

### SAβGal Histochemistry

Cells were washed twice with PBS then fixed with 15% Paraformaldehyde in PBS for 6 minutes. Cells were rinsed 3 times with PBS before staining with X-gal chromogenic substrate, which is cleaved by endogenous Beta galactosidase. Plates were kept in the staining solution, Parafilmed, to prevent from drying out, and incubated overnight at 37°C with ambient CO_2_. The next day, cells were washed again with PBSbefore switching to a 70% glycerol solution for imaging under a Leica brightfield microscope.

### Fixed and Live Cell Imaging

Samples were imaged using fluorescent microscopes – the Operetta High Content Imaging System (Perkin Elmer) or the BZ-X700 (Keyance) - and either a 10x or 20x air objective. Harmony (Operetta) or Volocity (BZ-X700) imaging software was used to adjust excitation and emission filters and came with pre-programmed AlexaFluor filter settings which were used whenever possible. All exposure times were optimized during the first round of imaging and then kept constant through all subsequent imaging.

### Image Analysis

Columbus (Operetta) or Image J (BZ-X700) was used for image analysis. Columbus software was to identify single cells utilizing DAPI of CellTracker Red to delineate nuclear and cell boundaries and calculate the signal statistics for each cell. Image J was used for muscle fibers to calculate the percentage of area composed of collagen by using the color threshold plug-in to create a mask of only the area positive for collagen. That area was then divided over the total area of the sample which was found using the free draw tool. All other fiber analyses were performed using Volocity software and manually counting fibers using the free draw tool.

### Statistical Analysis

All statistical analyses were performed using Matlab R2017a (MathWorks Software) or GraphPad Prism 5 (GraphPad Software). For statistical analysis, cellular or replicate distributions were compared by ttest before and after treatment of the same individual patient while aged vs young were compared after pooling distributions over three aged and three young samples run together (since of course the two cohorts were not patient matched); for ease of depiction the young, aged and treated boxplots show pooled distributions over all patient samples with the asterisks signifying the lowest significance level of all the paired comparisons, with one sample grace, given patient to patient variability.

### Cytokine Profiling

This work was performed together with the Human Immune Monitoring Center at Stanford University. Cell media was harvested and spun at 400 rcf for 10 minutes at room temperature. The supernatant was then snap frozen with liquid nitrogen until analysis. Analysis was done using the human 63-plex kit (eBiosciences/Affymetrix). Beads were added to a 96 well plate and washed in a Biotek ELx405 washer. Samples were added to the plate containing the mixed antibody-linked beads and incubated at room temperature for 1 hour followed by overnight incubation at 4°C with shaking. Cold and room temperature incubation steps were performed on an orbital shaker at 500-600 rpm. Following the overnight incubation, plates were washed in a Biotek ELx405 washer and then biotinylated detection antibody added for 75 minutes at room temperature with shaking. Plates were washed as above and streptavidin-PE was added. After incubation for 30 minutes at room temperature, wash was performed as above and reading buffer was added to the wells. Each sample was measured in duplicate. Plates were read using a Luminex 200 instrument with a lower bound of 50 beads per sample percytokine. Custom assay Control beads by Radix Biosolutions were added to all wells.

### Antibodies

The following antibodies were used in this study. The source of each antibody is indicated. Rabbit:: H3K9me3 (Abcam #ab8898 1: 4000), LAP2α (Abcam #ab5162 1:500), SIRT1 (Abcam #ab7343 1:200); Rabbit: Mouse: HP1γ (Millipore Sigma #05-690 1:200), Lamin A/C (Abcam #ab40567 1:200), GFP (Invitrogen, #A11122, 1:250); Luciferase (Sigma-Aldrich, #L0159, 1:200); Collagen I (Cedarlane Labs, #CL50151AP, 1:200); HSP47 (Abcam, #ab77609, 1:200), Laminin (Abcam, #AB11576, 1:1000), anti-CD31-Alexa Fluor 488 (clone WM59; BioLegend; #303110, 1:75), anti-CD45-Alexa Fluor 488 (clone HI30; Invitrogen; #MHCD4520, 1:75), anti-CD34-FITC (clone 581; BioLegend; #343503, 1:75), anti-CD29-APC (clone TS2/16; BioLegend; #303008, 1:75) and anti-NCAM-Biotin (clone HCD56; BioLegend; #318319, 1:75), anti-CD31-Alexa Fluor 488 (clone WM59; BioLegend; #303110, 1:75), anti-CD45-Alexa Fluor 488 (clone HI30; Invitrogen; #MHCD4520, 1:75), anti-CD34-FITC (clone 581; BioLegend; #343503, 1:75), anti-CD29-APC (clone TS2/16; BioLegend; #303008, 1:75) and anti-NCAM-iotin (clone HCD56; BioLegend; #318319, 1:75).

### RNA-Sequencing And Data Analysis

Cells were washed and digested by TRIzol (Thermo). Total RNA was isolated using the Total RNA Purification Kit (Norgen Biotek Corp) and RNA quality was assessed by the RNA analysis screentape (R6K screentape, Agilent); RNA with RIN > 9 was reverse transcribed to cDNA. cDNA libraries were prepared using 1 µg of total RNA using the TruSeq RNA Sample Preparation Kit v2 (Illumina). RNA quality was assessed by an Agilent Bioanalyzer 2100; RNA with RIN > 9 was reverse transcribed to cDNA. cDNA libraries were prepared using 500 ng of total RNA using the TruSeq RNA Sample Preparation Kit v2 (Illumina) with the added benefit of molecular indexing. Prior to any PCR amplification steps, all cDNA fragment ends were ligated at random to a pair of adapters containing a 8bp unique molecular index. The molecular indexed cDNA libraries were than PCR amplified (15 cycles) and then QC’ed using a Bioanalzyer and Qubit. Upon successful QC, they were sequenced on an Illumina Nextseq platform to obtain 80-bp single-end reads. The reads were trimmed by 2 nt on each end to remove low quality parts and improve mapping to the genome. The 78 nt reads that resulted were compressed by removing duplicates, while keeping track of how many times each sequence occurred in each sample in a database. The unique reads were then mapped to the human genome using exact matches. This misses reads that cross exon–exon boundaries, as well as reads with errors and SNPs/mutations, but it does not have substantial impact on estimating the levels of expression of each gene. Each mapped read was then assigned annotations from the underlying genome. In case of multiple annotations (e.g. a miRNA occurring in the intron of a gene), a hierarchy based on heuristics was used to give a unique identity to each read. This was then used to identify the reads belonging to each transcript and coverage over each position on the transcript was established. This coverage is non-uniform and spiky. Therefore, we used the median of this coverage as an estimate of the expression value of each gene. In order to compare the expression levels in different samples, quantile normalization was used. Further data analysis was done in Matlab. Ratios of expression levels were then calculated to estimate the log (base 2) of the fold-change. Student’s t-test was used to determine significance with a p<0.05 cutoff. Molecular Signatures Database categorization was done using Broad Institute online tools https://software.broadinstitute.org/gsea/msigdb/.

### Gene expression analysis

Total RNA was purified using RNeasy Plus Mini kit (Qiagen) and cDNA was prepared with First-strand cDNA synthesis kit (Applied Biosystems). The quantitative polymerase chain reaction (qPCR) was performed using VeriQuest Mastermix (Thermo Fisher Scientific) for SYBR Green and Taqman primer sets respectively. The relative gene expression was analyzed by the ΔΔCt method and normalized to glyceraldehyde-3-phosphate dehydrogenase (GAPDH). The expression levels were calculated using the method 2^-ΔΔCt. The Taqman probes for human GAPDH (Hs02758991); COL2A1 (Hs00264051); SOX9 (Hs00165814); MMP3 (Hs00233962) and MMP13 (Hs00233992) were purchased from Applied Biosystems. The SYBR green primer sequences used are: Human SOD2 (F)-5’GGC CTA CGT GAA CAA CCT GA3’; Human SOD2 (R)-5’TGG GCT GTA ACA TCT CCC TTG3’; Human iNOS (F)-5’GTC CCG AAG TTC TCA AGG CA3’; Human iNOS (R)-5’GTT CTT CAC TGT GGG GCT TG3’; Human RANKL (F)-5’CAG GTT GTC TGC AGC GT3’ and Human RANKL (R)-5’GAT CCA TCT GCG CTC TGA AAT A3’; Human GAPDH (F)- 5’TGT CCC CAC TGC CAA CGT GTC3’; Human GAPDH (R)-5’AGC GTC AAA GGT GGA GGA GTG GGT3’.

### ATP Assay

ATP in the chondrocytes was measured using colorimetric assay using ATP assay kit (ab83355; Abcam, Cambridge, MA) following manufacturer’s instructions. Cells were washed in cold phosphate buffered saline and homogenized and centrifuged to collect the supernatant. The samples were loaded with assay buffer in triplicate. ATP reaction mix and background control (50 µL) was added to the wells and incubated for 30?min in dark. The plate was read at OD 570 nm using SpectraMax M2e (Molecular Devices, Sunnyvale, CA). The mean optical density was used to estimate of the intracellular ATP concentration relative to the standard curve.

### Cell Proliferation Assay

Cell viability was assayed using the PrestoBlue Cell Viability (Life Technologies) reagent consecutively for 3 days post transfection in accordance to the manufacturer’s instructions. PrestoBlue reagent was added to the cell culture medium, and the cells were incubated at 30°C for 30min. Absorbance of the PrestoBlue was measured daily using SpectraMax M2e (Molecular Devices, Sunnyvale, CA).

### EDU staining

Staining was done according to the manufacter protocol using the Click-iT EdU kit. Cells were label with Edu after switching to growth media. Cells were allowed to grow one or two days before fixation with 4% paraformaldehyde and permeabilizationwith 0.5% Triton X-100 in PBST. Cells were the incubated in Click-It reaction cocktail for 30 min before washing in PBS and imaging.

### MitoTracker staining and flow cytometry analysis

Cells were washed twice with Ham’s F10 (no serum or pen/strep). Subsequently, MuSCs were stained with MitoTracker Green FM (ThermoFisher, M7514) and DAPI for 30 minutes at 37°C, washed three times with Ham’s F10, and analyzed using a BD FACSAria III flow cytometer.

### Myogenic colony-forming cell assay for MuSCs

Single treated and control MuSCs were deposited into wells of collagen- and laminin-coated plates at 1 cell per well by BD FACSAria III flow cytometer. Collagen/laminin coating was accomplished by overnight incubation of the plates rocking at 4°C with a 1:1 mixture of laminin (10 µg/ml ThermoFisher 23017-015) and collagen (10 µg/ml Sigma C8919) in PBS. Coated wells were washed three times with PBS before use. The cells were cultured in grow media, F10 medium supplemented with 20% horse serum and 5 ng/ml basic fibroblast growth factor (bFGF; PeproTech 100-18B). After 6 days of culture, plates were fixed with 4% paraformaldehyde (Electron Microscopy Services 15710), stained with DAPI (Invitrogen D1306), and scored by microscopy to determine the number of myogenic colony-forming cells (MCFCs), defined by wells that contained at least 8 cells.

### Myogenic/Fusion Index

Myogenic analysis was completed as previously described (16533505). After MuSCs underwent reprogramming or control treatment, cells were cultured in grow media. To induce differentiation, myoblast cultures were maintained in DMEM supplemented with 2% horse serum. The myogenic/fusion index was determined as the percentage of myonuclei in myotubes (defined as cells with three or more nuclei) compared with the total number of nuclei in the field.

### Lentiviral Transduction

Luciferase and GFP protein reporters were subcloned into a third generation HIV-1 lentiviral vector (CD51X DPS, SystemBio). To transduce freshly isolated MuSCs, cells were plated on a Poly-D-Lysine (Millipore Sigma, A-003-E) and ECM coated 8-well chamber slide (Millipore Sigma, PEZGS0896) and were incubated with 5 µl of concentrated virus per well and 8 µg/mL polybrene (Santa Cruz Biotechnology, sc-134220). Plates were spun for 5 min at 3200*g*, and for 1 hour at 2500*g* at 25**°**C. Cells were then washed with fresh media two times, scraped from plates, and resuspended in the final volume according to the experimental conditions.

### Bioluminescence Imaging

Bioluminescent imaging was performed using the Xenogen IVIS-Spectrum System (Caliper Life Sciences). Mice were anesthetized using 2% isoflurane at a flow rate of 2.5 l/min Intraperitoneal injection of D-Luciferin (50 mg/ml, Biosynth International Inc.) dissolved in sterile PBS was administered. Immediately following the injection, mice were imaged for 30 seconds at maximum sensitivity (f-stop 1) at the highest resolution (small binning). Every minute, a 30 second exposure was taken until the peak intensity of the bioluminescent signal began to diminish. Each image was saved for subsequent analysis.

### Bioluminescence Image Analysis

Analysis of each image was performed using Living Image Software, version 4.0 (Caliper Life Sciences). A manually-generated circle was placed on top of the region of interest and resized to completely surround the limb or the specified region on the recipient mouse. Similarly, a background region of interest was placed on a region of a mouse outside the transplanted leg.

### Tissue Harvesting

TA muscles were carefully dissected away from the bone, weighed, and placed into a 0.5% PFA solution for fixation overnight. The muscles were then moved to a 20% sucrose solution for 3 hours or until muscles reached their saturation point and began to sink. The tissues were then embedded and frozen in Optimal Cutting Temperature (OCT) medium and stored at −80°C until sectioning. Sectioning was performed on a Leica CM3050S cryostat that was set to generate 10 µm sections. Sections were mounted on Fisherbrand Colorfrost slides. These slides were stored at −20°C until immunohistochemistry could be performed.

### Histology

TA muscles were fixed for 5 hours using 0.5% electron-microscopy-grade paraformaldehyde and subsequently transferred to 20% sucrose overnight. Muscles were then frozen in OCT, cryosectioned at a thickness of 10 µm, and stained. For colorimetric staining with Hematoxylin and Eosin (Sigma) or Gomorri Trichrome (Richard-Allan Scientific), samples were processed according to the manufacturer’s recommended protocols.

### Ex vivo force measurement

To measure the force, we isolated the TA in a bath of oxygenated Ringer’s solution and stimulated it with plate electrodes. Immediately after euthanasia, the distal tendon of the TA, the TA, and the knee (proximal tibia, distal femur, patella, and associated soft tissues) were dissected out and placed in Ringer’s solution (Sigma) maintained at 25°C with bubbling oxygen with 5% carbon dioxide. The proximal tibia was sutured to a rigid wire attached to the force transducer and the distal tendon was sutured to a rigid fixture. No suture loops or slack was present in the system. The contralateral limb was immediately dissected and kept under low passive tension in oxygenated Ringer’s solution bath until measurement.

Supramaximal stimulation voltage was found and the active force-length curve was measured in a manner similar to the in vivo condition. After measurement, the muscle was dissected free and the mass measured. An Aurora Scientific 1300-A Whole Mouse Test System was used to gather force production data.

### DNA methylation data

The human Illumina Infinium EPIC 850K chip was applied to n=16 DNA samples (corresponding to 2 two treatment levels (before/after treatment) of 4 fibroblasts and 4 endothelial cells). The raw image data were normalized using the “preprocessQuantile” normalization method implemented in the “minfi” R package ^39,40^.

### Epigenetic clock analysis

Several DNAm based biomarkers have been proposed in the literature which differ in terms of their applicability (most were developed blood) and in terms of their biological interpretation (reviewed in ^15^). We focused on two epigenetic clocks that apply to fibroblasts and endothelial cells. In our primarly analysis, we used the pan-tissue epigenetic clock ^5^ because it applies to all sources of DNA (with the exception of sperm). A previously defined mathematical algorithm is used to combine the methylation levels of 353 CpG into an age estimate (in units of years), which is referred to as epigenetic age or DNAm age. In our secondary analysis, we used the skin & \blood epigenetic clock (based on 391 CpGs) because it is nown to lead to more accurate DNAm age estimates in fibroblasts, keratinocytes, buccal cells, blood cells, saliva, endothelial cells^17^.

We used the online version of the epigenetic clock software to arrive at DNA methylation age estimates from n=16 samples collected from n=8 individuals^5^. Although the chronological age range was relatively narrow (ranging from 47 years to 69 years, median age=55), the two DNAm age estimates exhibited moderately high correlations with chronological age (r=0.42 and r=0.63, P0.0089 for the pan tissue clock and the skin & blood clock, respectively).

Two samples (before and after rejuvenation treatment) were generated from each of n=8 individuals. To properly account for the dependence structure in the data, we used linear mixed effects models to regress DNAm age (dependent variable) on treatment status, chronological age, and individual identifier (coded as random effect). Toward this end, we used the “lmer” function in the “lmerTest” R package (Alexandra Kuznetsova, Per B. Brockhoff and Rune H. B. Christensen (2017) lmerTest Package: Tests in Linear Mixed Effects Models. Journal of Statistical Software, 82(13), 1–26. doi:10.18637/jss.v082.i13)

